# Synthesizing unmodified, supercoiled circular DNA molecules *in vitro*

**DOI:** 10.1101/2025.01.24.634800

**Authors:** Sepideh Rezaei, Monica Moncada-Restrepo, Sophia Leng, Jeremy W. Chambers, Fenfei Leng

**Author notes:** Current address: Department of Chemistry, Johns Hopkins University, 138 Remsen Hall, 3400 N. Charles Street, Baltimore, MD 21218. Current address: Department of Biology & Microbiology, Alfred Dairy Science Hall 228, Box 2104A, South Dakoda State University, Brookings, SD 57007. To whom correspondence should be addressed: Department of Chemistry & Biochemistry, Florida International University, 11200 SW 8th Street, FL 33199.

## Abstract

Supercoiled (Sc) circular DNA, such as plasmids, has shown therapeutic potential since the 1990s, but is limited by bacterial modifications, unnecessary DNA sequences, and contaminations that may trigger harmful responses. To overcome these challenges, we have developed two novel scalable biochemical methods to synthesize unmodified Sc circular DNA. Linear DNA with two loxP sites in the same orientation is generated via PCR or rolling circle amplification. Cre recombinase then converts this linear DNA into relaxed circular DNA. After T5 exonuclease removes unwanted linear DNA, topoisomerases are employed to generate Sc circular DNA. We have synthesized EGFP-FL, a 2,002 bp mini-circular DNA carrying essential EGFP expression elements. EGFP-FL transfected human HeLa and mouse C2C12 cells with much higher efficiency than *E. coli*-derived plasmids. These new biochemical methods can produce unmodified Sc circular DNA, in length from 196 base pairs to several kilobases and in quantities from micrograms to milligrams, providing a promising platform for diverse applications.

---

In 1990, Wolff and colleagues demonstrated that injection of a supercoiled (Sc) plasmid containing the firefly luciferase gene into mouse skeletal muscle resulted in the uptake and expression of the gene.^1, 2^ The luciferase activity was detectable in muscle for at least two months.^1^ Since then, plasmid DNA molecules have been extensively explored as therapeutics and have shown great potential.^3–6^ For example, ZyCov-D, a plasmid-based COVID-19 vaccine, has been approved for emergency use in India.^4, 7, 8^ Several plasmid-based vaccines are approved for veterinary use in animals.^9–12^ Plasmids are also widely used in gene therapy, with 12.6% of all gene therapy clinical trials (483 clinical trials) by 2023 utilizing plasmids (https://a873679.fmphost.com/fmi/webd/GTCT). These clinical trials have targeted various diseases including cystic fibrosis,^13^ cancers,^3, 14^ diabetes,^15, 16^ cardiovascular diseases,^17^ and _HIV._18-20

Currently, almost all plasmids are produced in *E. coli K-12* strains.^3, 21–23^ However, this production process has several drawbacks that pose potential risks for clinical applications. Plasmids isolated from *E. coli* typically carry modified bases, such as 6-methyladenine (6mA) in the 5’-GATC-3’ sequences (Dam sites) and 5-methylcytosine (5mC) in the 5’-CCWGG-3’ sequences (Dcm sites) that do not exist in human cells.^24–27^ Humans also lack enzymes, such as methylases and demethylases, to remove or modify these methylated bases. The potential health risks associated with these base modifications are not yet fully assessed and understood.^3, 28, 29^ Furthermore, due to the essential role of Dam methylation in *E. coli*, ^24, 30–32^ producing unmodified or unmethylated plasmids in *E. coli* for therapeutic use is currently not feasible.^33^

Another issue is that plasmids isolated from *E. coli* typically contain bacterial DNA sequences, such as a DNA replication origin and an antibiotic resistance-encoding gene, that are necessary for propagation and selection in *E. coli*.^21^ These DNA sequences increase the size of the plasmid and may trigger immune response and gene silencing.^34^ Additionally, antibiotic resistance genes pose the risk of horizontal transfer to bacteria in the human microbiome.^35, 36^ To address this issue, DNA minicircles—produced *in vivo* using site-specific recombination and consisting mainly of the target gene unit without bacterial DNA sequences—have been explored for clinical applications.^6, 37–41^ However, parent plasmid contamination remains a concern^42^ and the high cost of producing minicircles remains a challenge.^43, 44^

Moreover, it is extremely difficult to completely remove unwanted contaminants, such as *E. coli* genomic DNA, RNA, protein, and residual endotoxins and antibiotics, from the final plasmid product for different applications including therapeutic use.^6, 45–48^ These contaminants may trigger immune reactions in patients, further complicating their clinical application.^6, 45–48^

Clearly, a novel and effective method for producing unmodified Sc circular DNA molecules is urgently needed for therapeutic applications. To address this critical need, we have developed two innovative biochemical methods for *in vitro* production of unmodified Sc circular DNA molecules. These methods involve either polymerase chain reaction (PCR) using *Taq* DNA polymerase or rolling circle amplification (RCA) using ϕ29 DNA polymerase to generate linear DNA molecules containing two loxP sites oriented in the same direction. Cre DNA recombinase subsequently converts these linear DNA molecules into relaxed (Rx) circular DNA molecules. T5 exonuclease is used to remove any unwanted linear DNA. Finally, *E. coli* DNA gyrase or *variola* DNA topoisomerase I is employed to transform the Rx circular DNA molecules into Sc circular DNA molecules.

Using these two biochemical methods, we synthesized several unmodified Sc circular DNA molecules including EGFP-FL, a 2,002 bp mini-circular DNA molecule that carries only the essential elements for expressing enhanced green fluorescent protein (EGFP) in mammalian cells. Our results showed that the *in vitro*-synthesized unmodified EGFP-FL efficiently transfected human HeLa cells and mouse C2C12 myoblast cells. Its transfection efficiency is much higher than that of *E. coli*-derived and -modified circular DNA molecules. Additionally, we have synthesized two small Sc minicircles which are excellent tools to study supercoiling-induced DNA bendability, looping, and other DNA physical properties.

## Results and discussion

Figure 1 shows our strategy to produce unmodified Sc circular DNA molecules through a combination of *in vitro* DNA synthesis using polymerase chain reaction (PCR) or rolling circle amplification (RCA), formation of the Rx circular DNA molecules through site-specific recombination by Cre DNA recombinase,^49, 50^ and selective digestion of unwanted linear DNA by T5 exonuclease.^51–53^ The final unmodified Sc circular DNA molecules will be generated by two methods. Method 1 is using *E. coli* DNA gyrase to convert the Rx circular DNA molecules to Sc circular DNA molecules in the presence of ATP.^51^ Method 2 is using a eukaryotic DNA topoisomerase I, such as *variola* DNA topoisomerase I, in the presence of ethidium bromide, to convert Rx circular DNA molecules into (-) Sc circular DNA molecules (Figure 1). Rx circular DNA can be temporarily (+) supercoiled by a DNA intercalator, such as ethidium bromide.^54^ *Variola* DNA topoisomerase I can relax the temporarily (+) Sc circular DNA. After phenol extraction removing ethidium bromide, (-) Sc circular DNA molecules are produced.

**Figure 1.**
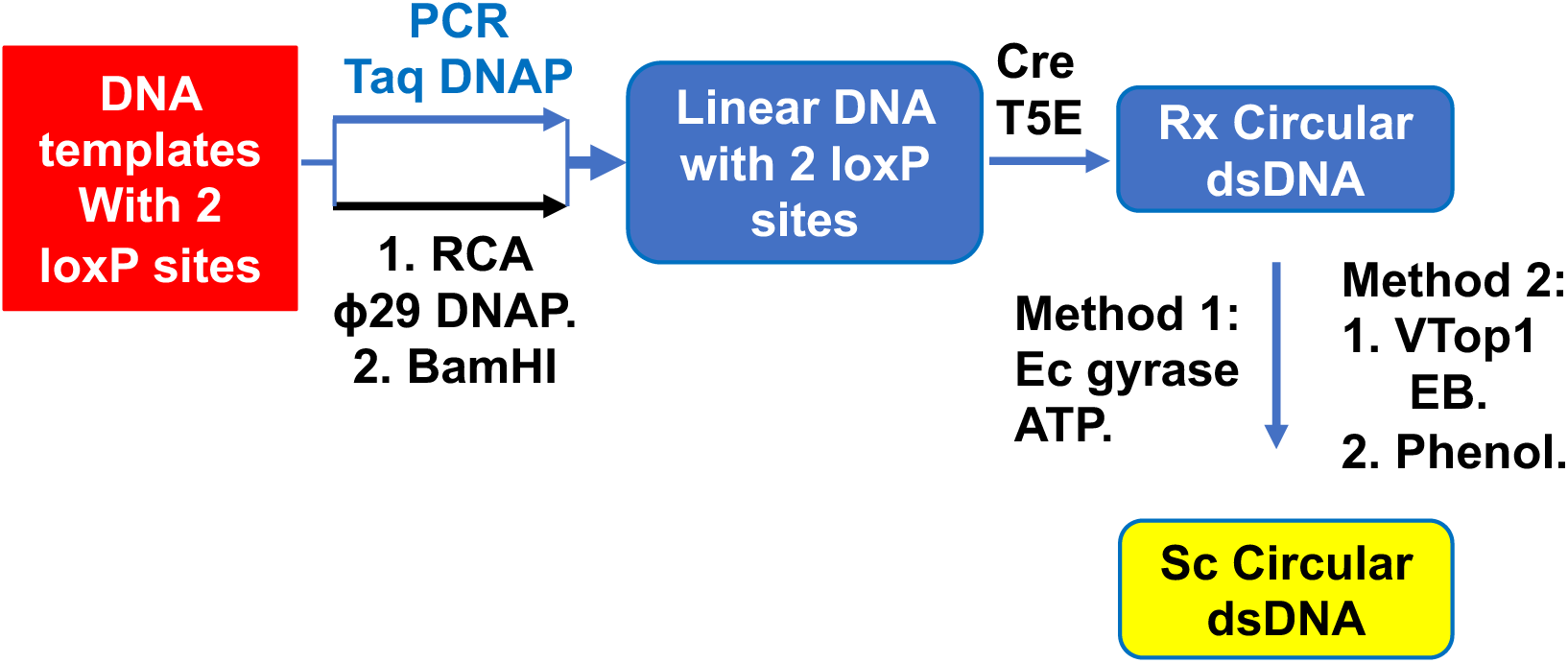
Strategies to synthesize Sc double-stranded circular DNA molecules. Symbols: PCR, polymerase chain reaction; DNAP, DNA polymerase; RCA, rolling circle amplification; Rx, relaxed; Sc, supercoiled; Cre, Cre recombinase; T5E, T5 exonuclease; EB, ethidium bromide; vTop1, variola virus DNA topoisomerase 1; Phenol, phenol extraction.

### A PCR-based biochemical method to synthesize unmodified, supercoiled double-stranded circular DNA molecules *in vitro*

We established a PCR-based biochemical method to synthesize unmodified Sc circular DNA molecules *in vitro* (Figure S1A). DNA templates with two loxP sites were used in the PCR reactions to produce double-stranded linear DNA molecules containing two loxP sites oriented in the same direction. Cre recombinase was employed to convert the linear DNA molecules into Rx circular DNA molecules.^49, 50^ T5 exonuclease was then used to digest unwanted linear DNA molecules.^51^ Subsequently, *E. coli* DNA gyrase or *variola* topoisomerase I was utilized to convert Rx circular DNA molecules to Sc circular DNA molecules. Figures 2 and 3 show two examples of the PCR-based biochemical method.

**Figure 2.**
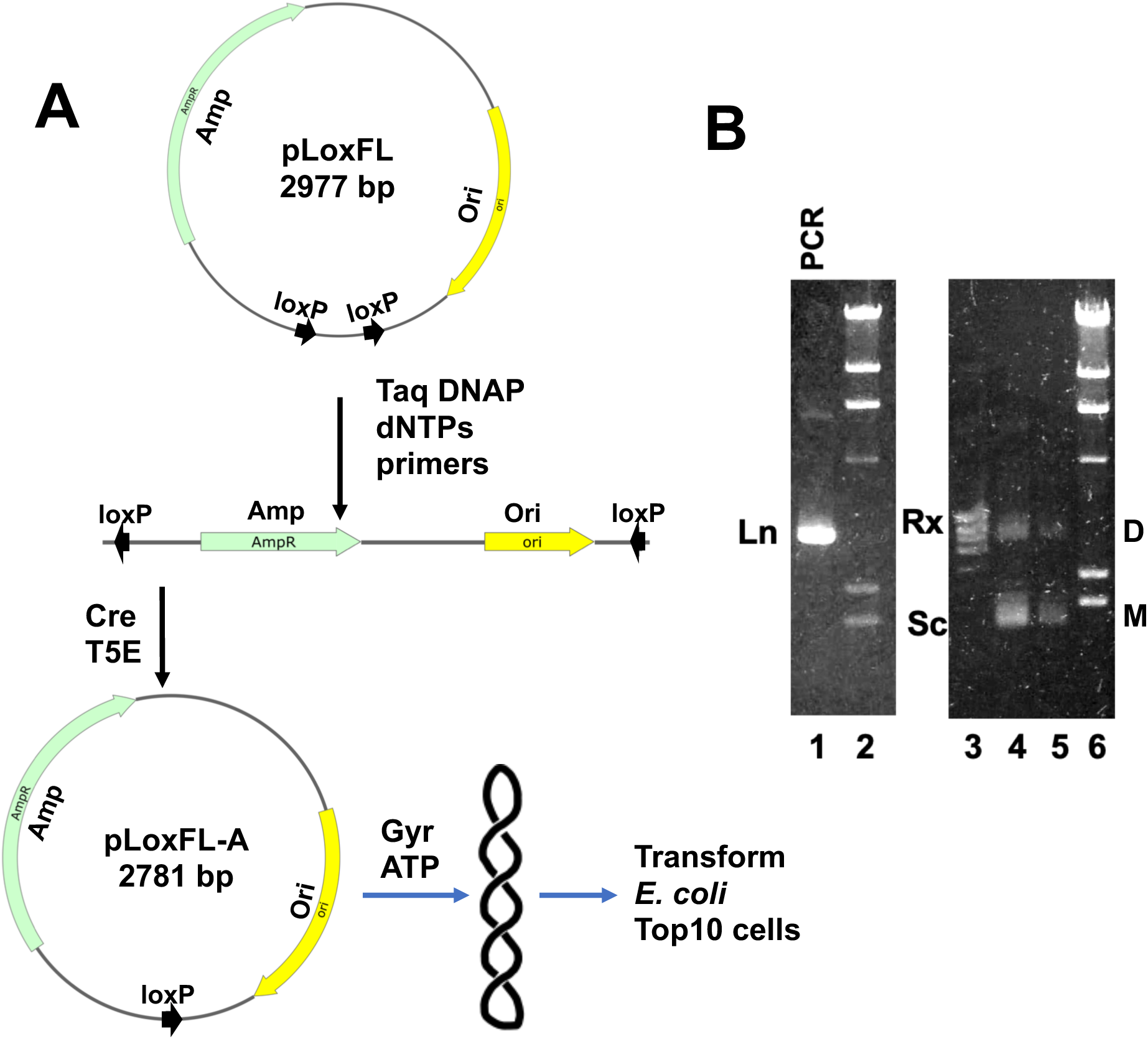
(**A**) Experimental procedure to generate Rx and Sc pLoxFL-A using the PCR-based method. (**B**) Lane 1 is the PCR product using FL1038 and FL1041 as primers and pLoxFL as the template. Lanes 2 and 6 are the λ DNA HindIII digest. Lane 3 is the relaxed (Rx) pLoxFL-A. Lanes 4 and 5 are the supercoiled (Sc) pLoxFL-A. Ln, linear; D, dimer; M, monomer. DNA sequencing confirmed the identity of the in vitro synthesized pLoxFL-A.

**Figure 3.**
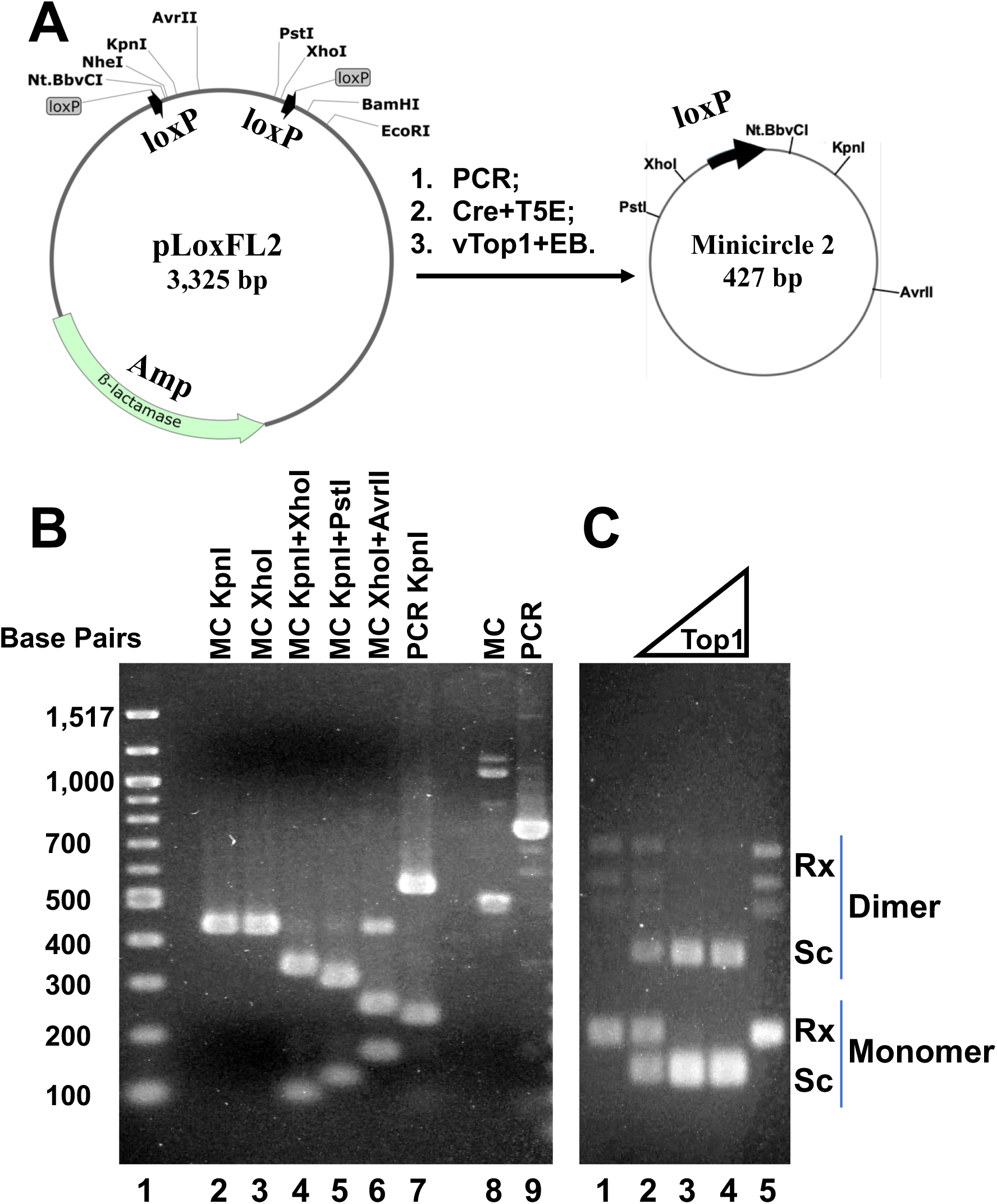
Synthesizing supercoiled (Sc) circular DNA molecule minicircle 2 using a PCR-based *in vitro* method. (**A**) Steps to synthesize Sc circular DNA molecule minicircle 2 (427 bp) using PCR and plasmid pLoxFL2 as the DNA template. Cre, Cre recombinase; T5E, T5 exonuclease; vTop1, variola DNA topoisomerase 1; EB, ethidium bromide. The identity of minicircle 2 was confirmed by DNA sequencing. (**B**) Minicircle 2 was digested by different restriction enzymes. MC, minicircle 2. (**C**) Minicircle 2 was negatively supercoiled by *variola* DNA topoisomerase 1 in the presence of 25 *μ*M EB. *E. coli* DNA gyrase could not completely supercoil minicircle 2.

Plasmid pLoxFL, a 2,977 bp plasmid isolated from *E. coli* cells, contains two loxP sites oriented in the same direction. A 2,870 bp linear PCR product carrying these two loxP sites (lane 1, Figure 2B) was generated through PCR using two specific primers targeting the smaller region of pLoxFL (Table S1; FL1064: 5′-CCATGCTGCAGGAATTCC-3′; FL1065: 5′-GACAGCTTATCATCGATAAGC-3′). Following recombination reactions with Cre recombinase and digestion with T5 exonuclease, a 2,781 bp Rx circular DNA molecule, pLoxFL-A, was produced (lane 3, Figure 2B). Sc pLoxFL-A was subsequently generated by treating pLoxFL-A with *E. coli* DNA gyrase in the presence of ATP (lanes 4 and 5, Figure 2B). Whole plasmid DNA sequencing confirmed the identity of the Rx and Sc circular DNA molecule pLoxFL-A. The Rx and Sc pLoxFL-A efficiently transformed *E. coli* Top10 cells. Again, DNA sequencing confirmed the identity of pLoxFL-A isolated from *E. coli* cells containing this plasmid.

Using a similar approach, a 427 bp minicircular DNA molecule (minicircle 2) was synthesized (Figure 3). PCR reactions generated a 779 bp linear DNA molecule containing two loxP sites oriented in the same direction (lane 9, Figure 3B; primers: FL1084F and FL1085R (Table S1)). A recombination reaction with Cre recombinase, followed by T5 exonuclease digestion, produced Rx minicircle 2 (lane 8, Figure 3B, and lane 1, Figure 3C). The identity of minicircle 2 was confirmed through restriction digestion assays (Figure 3B) and DNA sequencing.

Interestingly, under our experimental conditions, *E. coli* DNA gyrase in the presence of ATP could not completely supercoil minicircle 2 (Figure S2). In contrast, *variola* DNA topoisomerase I, when used in the presence of 25 µM ethidium bromide followed by phenol extraction, successfully and completely supercoiled minicircle 2 (Figure 3C). A likely explanation is that wrapping the 427 bp minicircle 2 around the large *E. coli* DNA gyrase imposes significant structural constraints on the small DNA minicircle (Figure S3). As a result, *E. coli* DNA gyrase cannot efficiently supercoil minicircle 2. On the other hand, *variola* DNA topoisomerase I, a smaller type IB topoisomerase, employs a controlled rotation mechanism to relax DNA molecules, which does not require wrapping the DNA around the enzyme.^55, 56^ The supercoiling efficiency is expected to be much higher (Figure 3C).

We also explored the possibility of including loxP sequences in both forward and reverse primers to amplify DNA sequences with two loxP sites. We reasoned that the 34 bp AT-rich loxP sequence, having a low melting temperature, would not significantly affect PCR reactions if incorporated into the 5′-ends of both forward and reverse primers (Figure S1B). If this hypothesis were correct, it would allow the production of linear DNA fragments carrying two loxP sites in the same orientation for any DNA sequence through PCR amplification (Figure S1). Rx and Sc circular DNA molecules could then be generated using Cre recombinase, T5 exonuclease, and *E. coli* DNA gyrase or a DNA topoisomerase (Figure S1B).

Our results, shown in Figures 4 and S4, confirmed that this PCR-based method can be used to produce Rx and Sc circular DNA molecules. Two primers with loxP sequences at their 5′-ends (FL1102F and FL1104R; Table S1) were used to amplify a large fragment of plasmid pGFPuv by PCR, yielding a 3,352 bp linear DNA fragment containing two loxP sites (lane 1, Figure 4B). Recombination using Cre recombinase, followed by T5 exonuclease digestion, produced a smaller Rx circular DNA molecule, plasmid pGFPuv-loxP (3,287 bp; lane 4, Figure 4B). DNA sequencing confirmed the identity of pGFPuv-loxP. The Rx pGFPuv efficiently transformed *E. coli* Top10 cells which were fluorescent under UV (Figure 4C). Again, DNA sequencing confirmed the identity of pGFPuv-loxP isolated from *E. coli* cells containing this plasmid.

**Figure 4.**
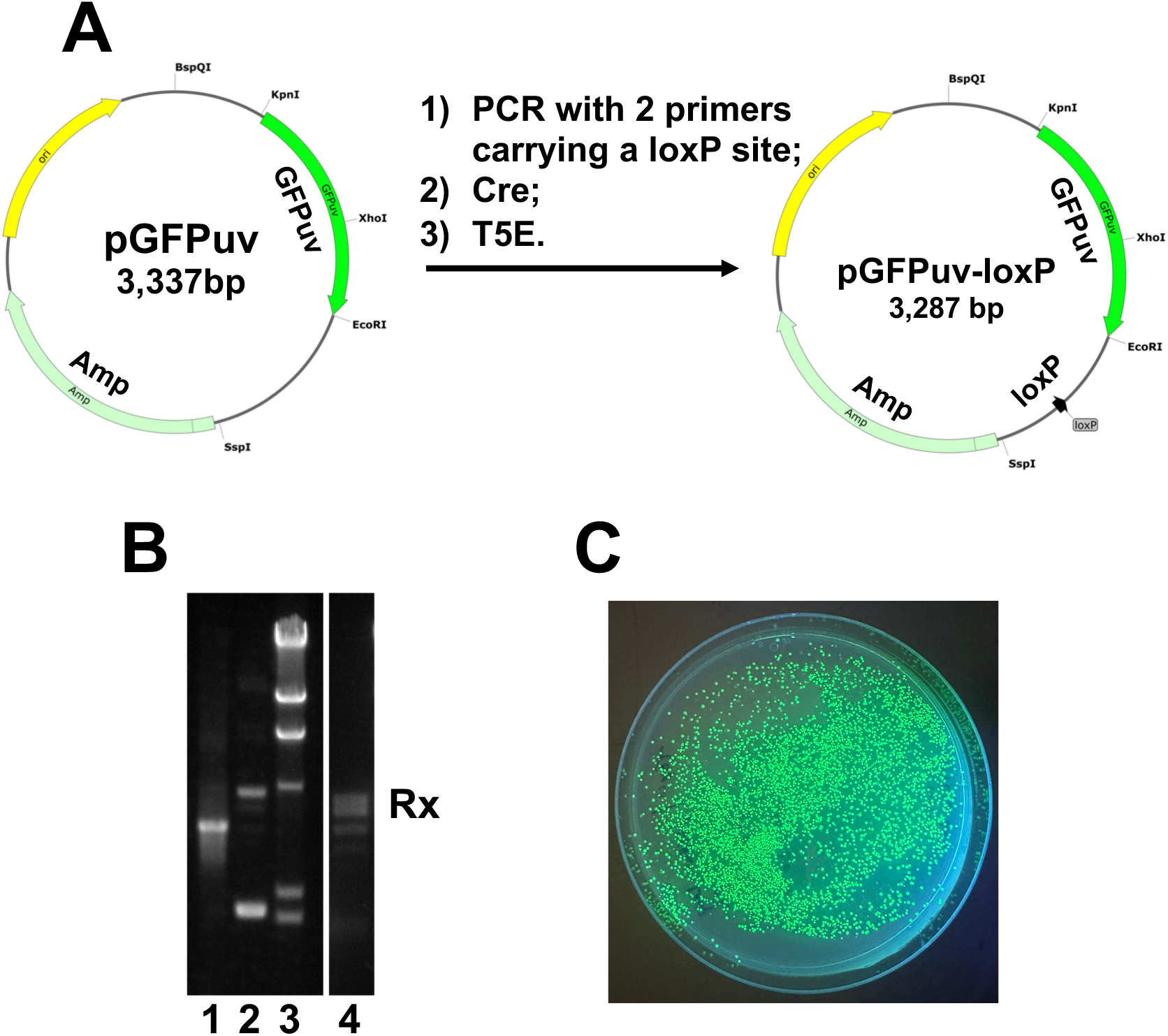
Circular DNA molecule pGFPuv-loxP was produced using plasmid pGFPuv as the DNA template that does not contain loxP sites by the PCR-based biochemical method (A, the procedure). Forward and reverse primers carrying a loxP site were used to amply a 3,352 bp DNA fragment of pGFPuv (B, lane 2) to generate a linear DNA fragment carrying two loxP sites facing the same orientation (B, lane 1). Cre DNA recombinase converted the linear DNA molecule into Rx pGFPuv-loxP. T5 exonuclease was used to digest unwanted linear DNA molecules (B, lane 4). (C) Fluorescence image of E. coli Top10 cells carrying pGFPuv-loxP.

Next, we synthesized a 1,933 bp unmodified minicircular DNA molecule, EGFP-FL1, using this PCR-based biochemical method. EGFP-FL1 contains only the essential DNA elements required for expressing EGFP in mammalian cells, including a CMV promoter and enhancer, an EGFP open reading frame, and an SV40 poly(A) signal (Figure S4). It does not include any additional sequences, such as an *E. coli* origin of replication. A PCR reaction using forward and reverse primers, each with a loxP site at the 5′-end, and a DNA template plasmid pEGFP-C1-FL, generated a 2,023 bp linear DNA fragment (lane 2, Figure S4B). Recombination using Cre recombinase, followed by T5 exonuclease digestion, produced the Rx form of EGFP-FL1 (lane 4, Figure S4). DNA sequencing confirmed the identity of EGFP-FL1. Due to the limited amount of Rx EGFP-FL1 produced, *E. coli* DNA gyrase was not used to convert the Rx form to the Sc form. Additionally, EGFP-FL1 was not used for transfection into mammalian cells. However, a similar minicircular DNA molecule, EGFP-FL, generated using an RCA-based method, was successfully converted to the Sc form and transfected into human HeLa cells and mouse C1C12 muscle cells, as detailed in the next section.

### An RCA-based biochemical method to synthesize unmodified, supercoiled double-stranded circular DNA molecules *in vitro*

We also established an RCA-based biochemical method to synthesize Sc circular DNA molecules by utilizing ϕ29 DNA polymerase (Figure 1). ϕ29 DNA polymerase^57^ from the Bacillus subtilis phage ϕ 29 is a DNA polymerase known for its high fidelity due to its inherent 3’→5’ proofreading exonuclease activity,^58^ extreme processivity,^59^ and exceptional strand displacement.^59^ Because ϕ29 DNA polymerase has these unique properties, it has been used for synthesizing large amount of single-stranded and double-stranded DNA,^60^ and also for whole genome amplification.^61, 62^ Here, we used this DNA polymerase to synthesize DNA molecules for the RCA-based biochemical method.

Figure 5A shows our procedure for synthesizing unmodified Sc pLoxFLA from plasmid pLoxFL using the RCA-based biochemical method. Our first step was optimizing experimental conditions for RCA reactions. We found that all circular DNA molecules—Sc, Rx, and Nk (nicked) DNA— could serve as templates for RCA reactions (Figure S5). However, Nk pLoxFL (lane 3, Figure 8B; pLoxFL nicked by Nt.BbvCI) produced significantly more RCA products (lane 4, Figure 8B; Table S2). For example, 30 ng of Nk pLoxFL used as the DNA template in a 50 µL RCA mixture containing 1×ϕ29 DNA polymerase buffer, ϕ29 DNA polymerase, and primers FL1038 and FL1041 typically yielded approximately 30 µg of high-molecular-weight double-stranded DNA after overnight amplification at 30 °C (∼15 hours). We determined that the optimal concentration of ϕ29 DNA polymerase for RCA reactions was 200–300 nM. Pentamer and hexamer random primers performed effectively in RCA-based reactions and were selected for subsequent experiments. Additionally, 0.5 mM dNTPs, an incubation temperature of 30 °C, and a duration of 15 hours were found to be optimal conditions for RCA reactions.

**Figure 5.**
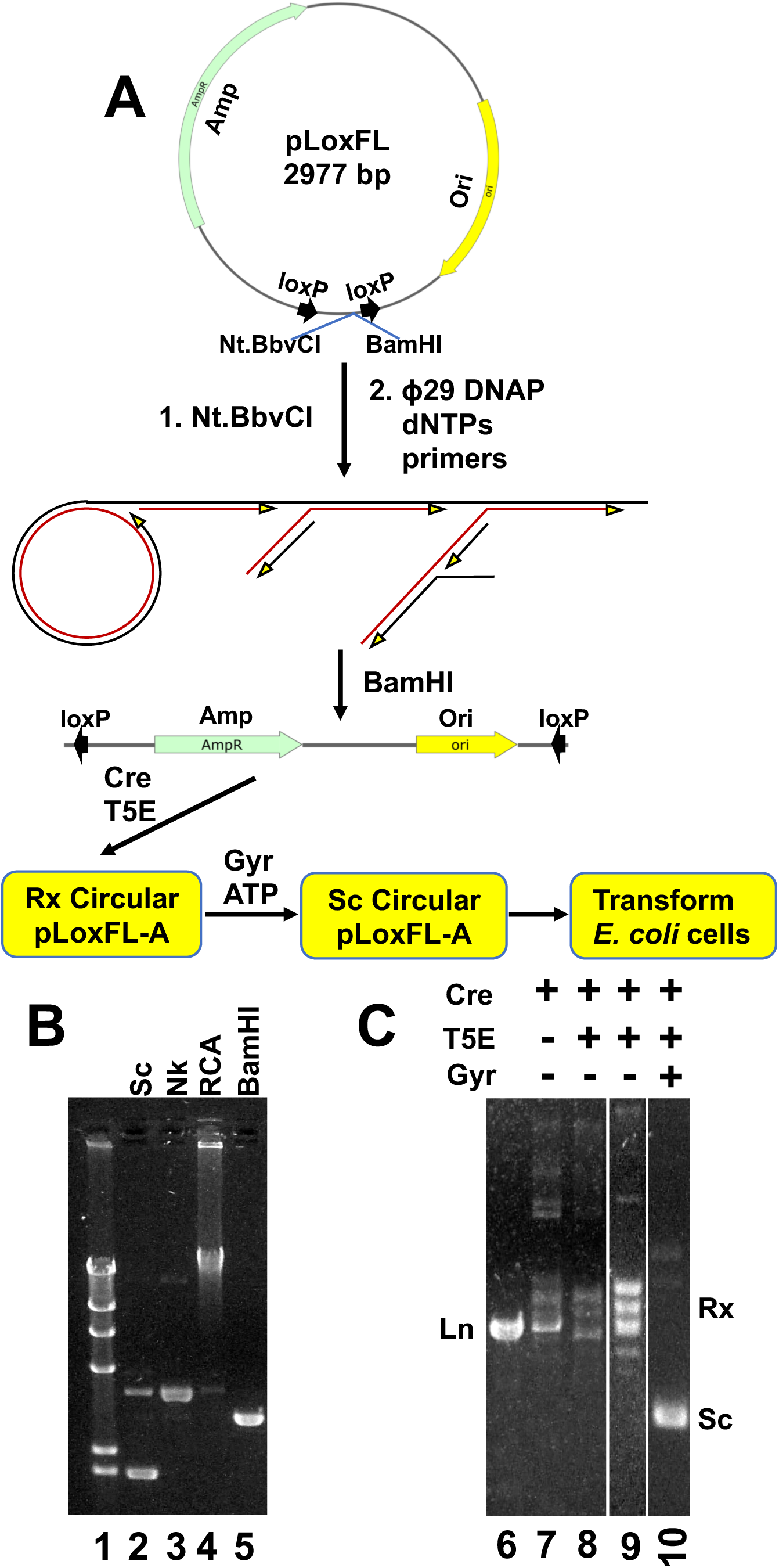
(A) Experimental procedure to generate Rx and Sc plasmid pLoxFL-A using the RCA-based method. (B) RCA of Nk pLoxFL by ϕ29 DNA polymerase. Lane 1, λ DNA HindIII digest. Lane 2, Sc pLoxFL. Lane 3, Nk pLoxFL. Lane 4, RCA product. Lane 5, the BamHI digest of the RCA product. (C) Generate Rx and Sc pLoxFL-A. Lane 6, RCA product BamHI digest. Lanes 7- 10, recombination products by Cre recombinase; lane 7, no T5 exonuclease; lanes 8-10, T5 exonuclease was added. Lane 9, purified and concentrated Rx pLoxFL-A. Lane 10, Sc pLoxFL- A by E. coli DNA gyrase.

After synthesizing sufficient amounts of high-molecular-weight double-stranded DNA using Nk pLoxFL as the template, BamHI digestion was performed to generate a linear 2.9 kb DNA fragment (Figure 8B, lane 5). The resulting linear DNA fragment, carrying two loxP sites (pLoxFL BamHI digest, lane 1, Figure 8C), was then converted into the Rx circular DNA molecule, pLoxFLA (lanes 8 and 9, Figure 8C), using Cre recombinase purified in our lab. The recombination efficiency was estimated to be approximately 74%, consistent with previous results.^49, 50^ T5 exonuclease completely removed unwanted linear and nicked DNA molecules (lane 9, Figure 8). Subsequently, *E. coli* DNA gyrase efficiently converted Rx pLoxFLA into Sc pLoxFLA (compare lanes 9 and 10, Figure 8). The final product also contained a small amount of Sc circular DNA dimers (lanes 9 and 10, Figure 8). DNA sequencing confirmed the identity of the unmodified Rx and Sc pLoxFLA.

Next, we synthesized unmodified Sc EGFP-FL, a 2,002 bp Sc circular DNA molecule, using the RCA-based biochemical method to study its transfection efficiency in mammalian cells (Figures 6A and 6B). EGFP-FL contains only the essential DNA elements required for expressing enhanced green fluorescent protein (EGFP) in mammalian cells, including a CMV promoter and enhancer, an EGFP open reading frame, and an SV40 poly(A) signal (Figure 6A). It does not include a bacterial origin of replication or any antibiotic resistance-encoding genes for selection and, therefore, cannot be replicated or produced in *E. coli*. The only bacterial DNA sequence present is the 34 bp loxP site (Figure 6A). Since the RCA-synthesized Sc EGFP-FL lacks bacterial modifications, such as Dam methylation, it was completely digested by DpnII and resistant to digestion by DpnI (Figure S6C). In contrast, Sc EGFP-FL generated from plasmid pLoxFL3, isolated from *E. coli* Top10 cells, contained Dam methylation sites. As a result, it was completely digested by DpnI but not by DpnII. This difference arises because both DpnI and DpnII recognize the same 5′-GATC-3′ Dam sequence: DpnI digests methylated Dam sequences, while DpnII digests unmethylated Dam sequences.^63^

**Figure 6.**
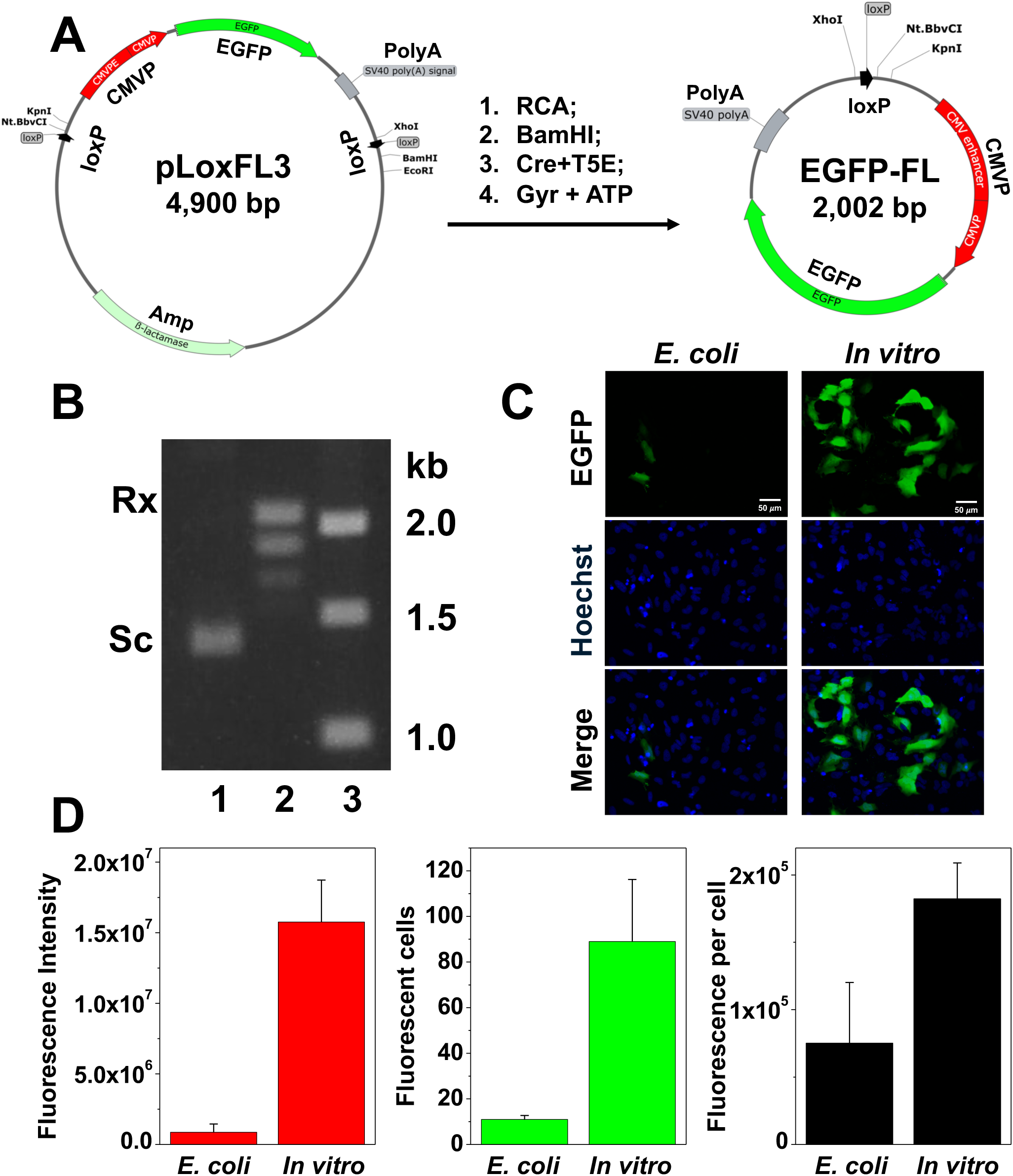
(**A**) Synthesizing Sc EGFP-FL using the RCA-based biochemical method. (**B**) 1% Agarose gel of Rx and Sc EGFP-FL. Lane 1, Sc EGFP-FL. Lane 2, Rx EGFP-FL. Lane 3, 1 kb DNA ladder. (**C**) The *in vitro*-synthesized, unmodified Sc EGFP-FL efficiently transfected human Hela cells. Hela cells were seeded and grown in 96-well plates in DMEM supplemented with 10% FBS for 24 hours. Subsequently, the cells were transfected, respectively, with 0.4 µg of EGFP-FL prepared from plasmid pLoxFL3 isolated from *E. coli* Top10 cells (left, *E. coli*) or *in vitro*- synthesized, unmodified EGFP-FL (right, *in vitro*) with PolyFect transfection reagent (Qiagen). Fluorescence images were captured at 48-hours post-transfection with a BZX800 fluorescence microscope. (**D**) The EGFP fluorescence intensity of transfected Hela cells by *E. coli* derived EGFP-FL or by *in vitro*-synthesized, unmodified EGFP-FL. Please note that the transfection efficiency of in vitro-synthesized, unmodified EGFP-FL was significantly higher than that using *E. coli* derived EGFP-FL.

Using the RCA-synthesized Sc EGFP-FL, we transfected human HeLa cells and mouse C2C12 myoblast cells. To our surprise, not only did the RCA-synthesized Sc EGFP-FL efficiently transfect both HeLa cells (Figures 6 and S7) and mouse C2C12 myoblast cells (Figure S8), but its transfection efficiency was substantially higher than that of Sc DNA molecules derived from *E. coli* Top10 cells. For example, we compared the transfection efficiency of the RCA-synthesized Sc EGFP-FL with the same Sc EGFP-FL generated using plasmid pLoxFL3 isolated from *E. coli*. The RCA-synthesized Sc EGFP-FL demonstrated significantly greater transfection efficiency in HeLa cells than the *E. coli*-derived Sc EGFP-FL (Figures 6C and 6D). Under our experimental conditions, the number of EGFP-positive cells and the fluorescence intensity per EGFP-positive cell were both markedly higher for the RCA-synthesized Sc EGFP-FL compared to the *E. coli*- derived counterpart (Figures 6C and 6D).

Similar results were observed when comparing the transfection efficiency of the RCA-synthesized Sc EGFP-FL with that of plasmid pEGFP-C1 isolated from *E. coli* Top10 cells in both human HeLa cells and mouse C2C12 cells (Figures S7 and S8). It is worth noting that increasing the excitation light intensity, exposure time, or gain allowed for the detection of more plasmid pEGFP- C1-transfected HeLa and C2C12 cells.

Furthermore, we synthesized a 196 bp minicircle, i.e., minicircle 4, using the RCA-based biochemical assay with Nk pLoxFL as the DNA template (Figure S9). RCA reactions, followed by PvuII digestion, produced a 2,977 bp linear DNA molecule containing two loxP sites oriented in the same direction (lane 2, Figure S9). Subsequent recombination with Cre recombinase and digestion with T5 exonuclease generated Rx minicircle 4 (lane 4, Figure S9). *Variola* DNA topoisomerase I in the presence of 25 µM ethidium bromide, followed by phenol extraction, produced Sc minicircle 4 (lane 5, Figure S9). Interestingly, the majority of minicircle 4 molecules were dimers (lanes 4 and 5, Figure S9), with some existing as monomers and tetramers. This is consistent with the small size of minicircle 4 and the DNA persistence length, which ranges between 100–150 bp in the presence of Mg^2+^ ^64, 65^. The high bending force required to form a 196 bp DNA minicircle likely contributed to the low yield of monomeric minicircle 4. Restriction enzyme digestion with BamHI, HindIII, and a combination of BamHI and HindIII produced DNA fragments consistent with the predicted lengths based on the minicircle 4 DNA map/sequence. Minicircles 2 and 4 are among the smallest DNA minicircles which can be used to study DNA biochemical and biophysical properties (ref).

### Advantages and potential applications of the *in vitro* synthesized, unmodified Sc circular DNA molecules

The unmodified Sc circular DNA molecules synthesized *in vitro* using the technology developed here offer several advantages over bacterial plasmid DNA and are expected to have potential applications in various fields, including use as therapeutics such as DNA vaccines and gene therapy.

First, the *in vitro*-synthesized Sc circular DNA molecules do not contain modified bases, making them less likely to be recognized as alien DNA and more readily accepted by human cells. In other words, they represent a much better choice for therapeutic applications compared to bacterial plasmid DNA. To support this, our results demonstrated that the *in vitro*-synthesized, unmodified Sc EGFP-FL efficiently transfected human HeLa cells and mouse C2C12 muscle cells (Figures 6, S7, and S8). The transfection efficiency of the *in vitro*-synthesized Sc EGFP-FL was significantly higher than that of bacterial-derived Sc circular DNA molecules (Figures 6, S7, and S8). One possible explanation for this difference is that bacterial plasmid DNA molecules contain methylated bases, including 6mA in 5’-GATC-3’ Dam sequences and 5mC in 5’-CCWGG-3’ Dcm sequences. Since human cells lack the specific methylases or demethylases for 6mA and also for 5mC in the Dcm sequences,^66^ these methylated bases may interfere with transfection efficiency and inhibit EGFP expression. Further studies are needed to elucidate the impact and mechanism of methylated bases on transfection efficiency and gene expression in mammalian cells.

Secondly, the *in vitro*-synthesized unmodified Sc circular DNA molecules, such as Sc EGFP-FL, do not contain unnecessary DNA sequences, such as bacterial DNA replication origins and antibiotic resistance genes, with the exception of a 34 bp loxP sequence. Consequently, these molecules are significantly smaller than their bacterial plasmid counterparts, which require a DNA replication origin and an antibiotic resistance gene for propagation and selection in *E. coli*. These extra sequence elements in bacterial plasmid DNA may also trigger immune responses and gene silencing, complicating therapeutic applications.^34^

Third, the *in vitro*-synthesized unmodified Sc circular DNA molecules are free from bacterial genomic DNA, RNA, endotoxins, and antibiotic contaminations. Since these two *in vitro* biochemical methods do not produce genomic DNA and RNA per se, contamination by genomic DNA and RNA is unlikely. However, trace amounts of genomic DNA may occasionally be introduced when Nk bacterial plasmids are used as DNA templates in PCR and RCA reactions or when purified proteins/enzymes are employed. These genomic DNA and Nk plasmid DNA contaminants can be effectively removed using DpnI digestion followed by T5 exonuclease treatment.^52, 63^

Fourth, these *in vitro* biochemical methods are scalable, capable of producing Sc circular DNA molecules in quantities ranging from micrograms to milligrams, grams, or even larger amounts. Enzymes such as ϕ29 DNA polymerase, Taq DNA polymerase, Cre recombinase, T5 exonuclease, *E. coli* DNA gyrase, and restriction enzymes (e.g., BamHI and EcoRI) can be overexpressed in *E. coli* and purified inexpensively. As a result, the production cost of *in vitro*-synthesized Sc circular DNA is expected to be comparable to that of plasmids isolated from *E. coli*. Additionally, the RCA-based method can be used to generate substantial quantities of linear DNA molecules, which can serve as templates for producing RNA molecules for various applications (Figure S9). Typically, 3-4 mg of linear DNA molecules can be produced using the procedure illustrated in Figure S9 for a 10 mL of reaction mixture (data not shown). Furthermore, these biochemical methods can be fully automated, offering a significant advantage over the plasmid manufacturing method using *E. coli* fermentation.

Fifth, the minicircles generated in this study serve as excellent tools and materials for investigating supercoiling-induced DNA bendability, looping, and other physical properties using single-molecule techniques, such as atomic force microscopy and cryo-EM, and DNA sequencing (data not shown).

Finally Taq DNA polymerase and ϕ29 DNA polymerase can use modified nucleotides for PCR^67^ or RCA^68–70^ reactions, enabling the production of Sc circular DNA molecules with modified nucleotides specific for various applications.

### Summary

In this research article, we present two novel *in vitro* biochemical methods for synthesizing Sc circular DNA molecules. These methods utilize either PCR or RCA to generate linear DNA molecules carrying two loxP sites oriented in the same direction. In the PCR-based method, the linear DNA molecules carrying two loxP sites can also be produced using forward and reverse primers, each containing a loxP site. Cre-mediated DNA recombination efficiently converts these linear DNA molecules into Rx circular DNA molecules. T5 exonuclease is then used to digest unwanted linear DNA. *E. coli* DNA gyrase with ATP or variola DNA topoisomerase I in the presence of ethidium bromide are employed to convert the Rx circular DNA molecules into Sc circular DNA molecules.

To demonstrate feasibility, we synthesized six different circular DNA molecules *in vitro* ranging from 196 base pairs to several kilobases in length: pLoxFLA, minicircle 2, pGFPuv-loxP, EGFP-FL, EGFP-FL1, and minicircle 4. Using EGFP-FL, a 2,002 bp Sc circular DNA molecule containing only the essential elements for expressing enhanced green fluorescent protein (EGFP) in mammalian cells, including a CMV promoter and an enhancer, an EGFP open reading frame, and an SV40 poly(A) signal, we tested the transfection efficiency of this Sc circular DNA molecule in HeLa cells and mouse C2C12 myoblast cells. The *in vitro*-synthesized Sc EGFP-FL not only transfected these cells efficiently but also exhibited significantly higher transfection efficiency compared to Sc DNA molecules derived from *E. coli*.

Unlike bacteria-derived plasmid DNA, the *in vitro*-synthesized circular DNA molecules are free of modified bases, unnecessary sequences (such as bacterial replication origins and antibiotic resistance genes), bacterial genomic DNA, RNA, endotoxins, and antibiotic contaminants. These features represent a significant advantage for various applications. Moreover, the scalability of these methods enables the production of Sc circular DNA molecules in quantities ranging from micrograms to potentially grams or more, making them suitable for use in therapeutics such as DNA vaccines and gene therapy and also for biochemical and biophysical studies. Furthermore, minicircles produced using these biochemical methods are excellent tools and materials for studying certain DNA biophysical properties.

## Methods

### Plasmids and oligonucleotides

Plasmid pLoxFL was constructed by inserting a 29 bp oligomer carrying BamHI and Nt.BbvCI sites into the PstI and AatII sites of pLOX2+.^71^ Plasmid pLoxFL2 was constructed by inserting a 690 bp synthetic DNA fragment carrying two loxP sites in the same direction into EcoRI and HindIII sites of pUC18. Plasmid pEGFP-C1-FL was constructed in two steps. First, plasmid pEGFP-C1^72^ was digested by using BamHI and BglII to remove the multiple cloning site and purified using Qiagen gel purification kit. The linearized DNA fragment was then religated by T4 DNA ligase to generate pEGFP-C1-FL. Plasmid pLoxFL3 was constructed by inserting a 1,903 bp PCR fragment carrying an EGFP gene and all other necessary components to express EGFP in mammalian cells amplified from pEGFP-C1-FL into the KpnI and XhoI sites of pLoxFL2. Plasmids pET28a(+)_His_Tev_Cre_recombinase, which expresses recombinant Cre DNA recombinase, and pET28a(+)_His_Tev_Phi29_DNAP, which express recombinant ϕ29 DNA polymerase, were synthesized and purchased from Gene Universal (https://www.geneuniversal.com/). All synthetic oligonucleotides including random hexamer primers and random pentamer primers were purchased from Eurofins Genomics, Inc. (+) Sc, (-) Sc, Nk, and Rx pLoxFL DNA samples were prepared as described previously.^51, 73^

### Proteins and enzymes

T5 exonuclease, *E. coli* DNA gyrase, and *variola* virus DNA topoisomerase I were purified as described previously.^51^ A His-tagged Cre DNA recombinase was purified from *E. coli* strain BLR(DE3) carrying plasmid pET28a(+)_His_Tev_Cre_recombinase by Ni-NTA column. The His-tag was then removed by TEV protease. A His-tagged ϕ29 DNA polymerase was purified from *E. coli* strain BLR(DE3) carrying plasmid pET28a(+)_His_Tev_Phi29_DNAP by Ni-NTA column followed by a SP-Sepharose Fast Flow column. The His-tag was removed by TEV protease as well. Thermal stable Taq DNA polymerase and SYBR Green were purchased from Fisher Scientific, Inc. Restriction enzymes Nt.BbvCI, BamHI, and EcoRI were purchased from New England Biolabs, Inc.

### Polymerase chain reaction (PCR)

PCR reactions were performed in a MJ Research PTC-200 thermal cycler using Taq DNA polymerase (1.5 units per 50 μL; Fisher Scientific, Inc), 0.2 mM of dNTPs, 0.5 μM of forward and reverse primers, and a DNA template in 1×PCR buffer (10 mM Tri-HCl, pH 8.8, 50 mM KCl, 1.5 mM MgCl_2_, and 0.08% NP-40). PCR thermal cycling conditions are: an initial denaturation step for 3 min at 95 °C, 27 cycles of denaturation for 30 seconds at 95 °C, annealing at 55 °C for 30 seconds, and extension at 72 °C for required amount of time (∼1 min per kb), and a final extension step at 72 °C for 10 min. PCR products were purified using GeneJET PCR Purification Kit (ThermoFisher Scientific, Inc) and analyzed in a 1% agarose gel by using 1×TAE buffer, pH 7.8.

### Rolling circle amplification (RCA) reactions by ϕ29 DNA polymerase

RCA reactions were performed using ϕ29 DNA polymerase, 0.5 mM dNTPs, 30 ng per 50 μL Nk circular DNA template or other circular DNA templates, 0.2 μM of a pair of specific primers or 5 μM of random hexamer or pentamer primers in 1×RCA reaction buffer (50 mM Tris-HCl, pH 7.5, 10 mM MgCl_2_, 10 mM (NH_4_)_2_SO_4_, 4 mM DTT, and 0.5 mg/mL of BSA) at 30 °C overnight or 14-16 hours. The high molecular weight DNA products of the RCA reactions may be digested by a restriction enzyme, such as BamHI or EcoRI, and then analyzed in in a 1% agarose gel by using 1×TAE buffer, pH 7.8.

### Generating circular DNA molecules by Cre recombinase

Recombination reactions by Cre recombinase (75 nM) were performed in 1×recombination buffer (50 mM Tris-HCl, pH 7.5, 33 mM NaCl, 10 mM MgCl_2_, and 1 mM DTT) at 37 °C for 30 min using a linear DNA template carrying two loxP sites in the same direction, which generates Rx circular DNA molecules and certain unwanted linear DNA molecules. The reaction mixtures were incubated at 65 °C for 10 min to inactivate Cre recombinase. Subsequently, the 0.2 μM of T5 exonuclease was added to the reaction mixtures and incubated at 37 °C for additional 2 hours to digest unwanted linear DNA molecules. The Rx circular DNA molecules were purified by using GeneJET PCR Purification Kit (ThermoFisher Scientific, Inc) or phenol extraction followed by ethanol or isopropanol precipitation and analyzed in in a 1% or 2% agarose gel by using 1×TAE buffer, pH 7.8.

### Cell culture

HeLa cells and mouse C2C12 cells were cultured in Dulbecco’s Modified Eagle Medium (DMEM) high glucose supplemented with 10% heat inactivated fetal bovine serum, 100 U/mL penicillin, and 100 µg/mL streptomycin (growth medium) in a humidified incubator at 37 °C and 5% CO2.

### Transfection of circular DNA containing EGFP into HeLa and C2C12 cells

HeLa cells and mouse C2C12 cells were grown in black, optical-bottom polystyrene 96-well plates (Thermo Scientific™ Nunc). HeLa and C2C12 cells were seeded at a density of 15,000 and 4,000 cells per well, respectively, in growth medium and incubated for 24 hours at 37 °C and 5% CO_2_. Subsequently, the cells were transfected with 0.4 µg of pEGFP-C1 isolated from *E. coli*, *E. coli*-derived EGFP-FL, or *in vitro*-synthesized unmodified EGFP-FL, using 1µL of PolyFect transfection reagent (Qiagen), according to the manufacturer’s instructions.

At 24- and 48-hours post-transfection, cells were incubated with 2 µg/mL Hoechst 33342 at room temperature for 30 minutes, prior to image acquisition. Fluorescent images were captured using identical acquisition parameters on either a Nikon Eclipse Ti-U Inverted Fluorescence Microscope or a Keyence BZX800 Fluorescence Microscope. Image analysis was performed using Fiji (National Institutes of Health) or BZ-X800LE Analyzer (Keyence) software. Total fluorescence intensity was quantified by measuring the integrated density (Fiji) or brightness (BZ-X800LE Analyzer) of the fluorescent cells, with consistent threshold settings applied across all images.

### Data availability

The data supporting the findings of this study are available within the paper and its Supplementary Information.

## Acknowledgments

This work was supported by National Institutes of Health grants 1R21AI125973, 1R21AI178134, and 1R41TR005250 (to F.L.).

## Author contributions

F.L. conceptualized the project and supervised the study. S.R.and S.L. performed most of the biochemical experiments and data processing. M.M.-R performed the cell biology studies. F.L. and J.C. discussed, analyzed, and interpret the data. F.L wrote the manuscript.

## Supplementary Information

Supplementary Information is available at *Nature Chemical Biology* Online.

