## Supplemental Figures and Tables for "Synthesizing unmodified, supercoiled circular DNA molecules *in vitro*"

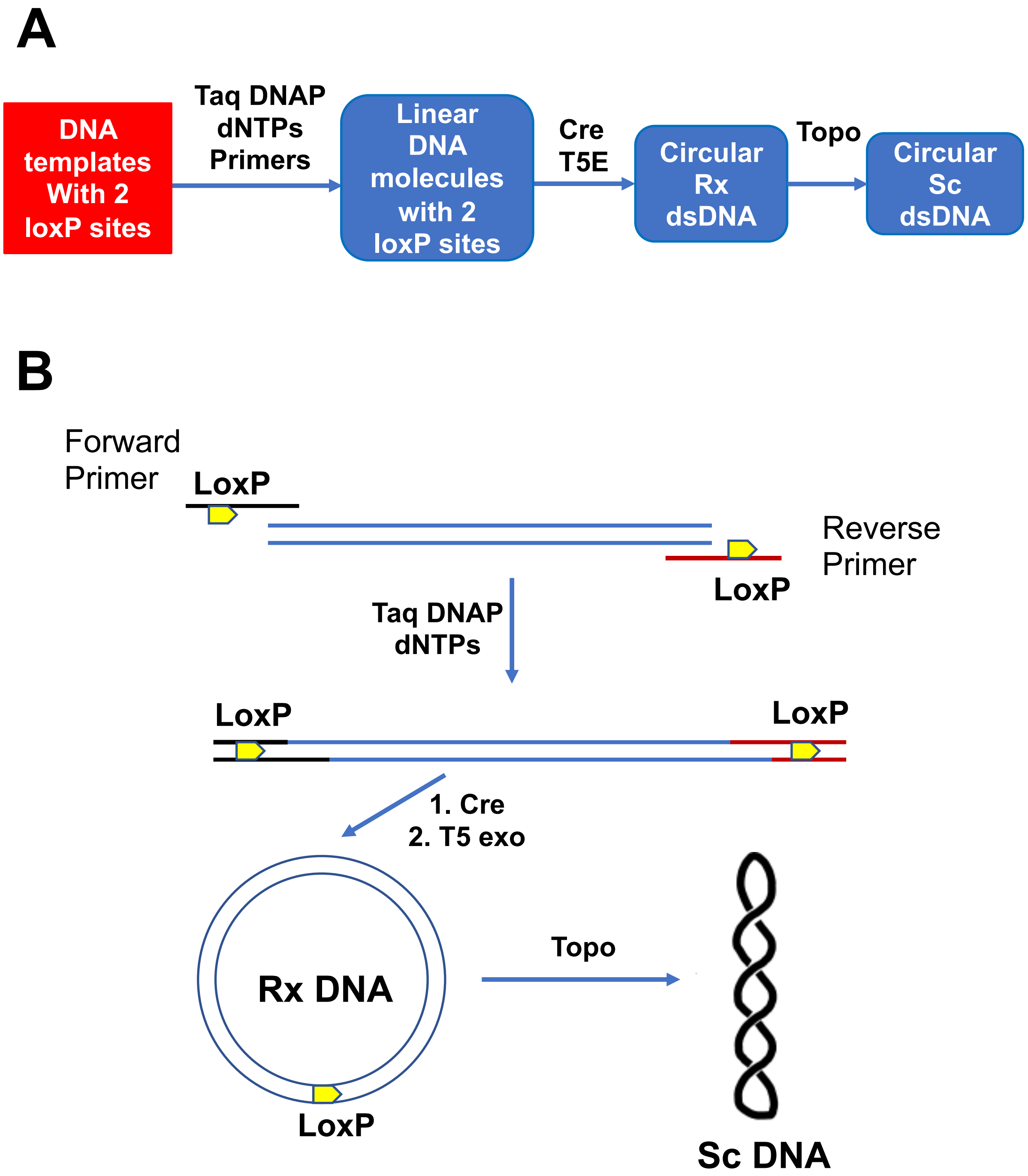


**Figure S1.** Strategies to synthesize Sc double-stranded circular DNA by PCR-based biochemical method. (**A**) DNA template carries two loxP site. (**B**) DNA templates do not contain loxP sites. LoxP sites are added to forward and reverse primers. DNAP, DNA polymerase; Rx, relaxed; Sc, supercoiled; Cre, Cre recombinase; T5E, T5 exonuclease; Topo, DNA topoisomerase.

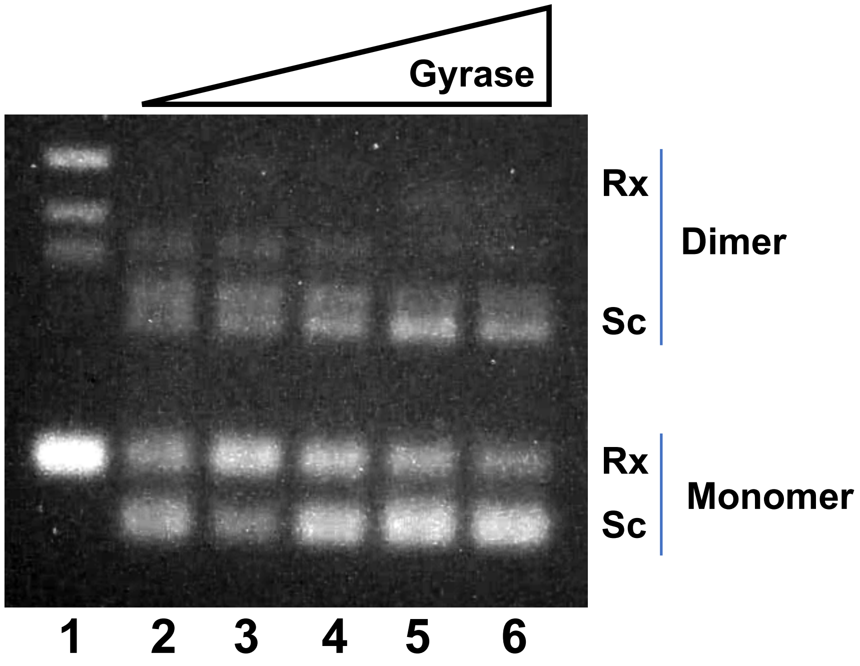


**Figure S2.** *E. coli* DNA gyrase could not completely supercoil Minicircle 2. DNA gyrase assays were described in Materials and Methods. Lane 1 is the relaxed Minicircle 2. Lanes 2-6 contain 50 nM of *E. coli* DNA gyrase, respectively. Rx, relaxed. Sc, supercoiled.

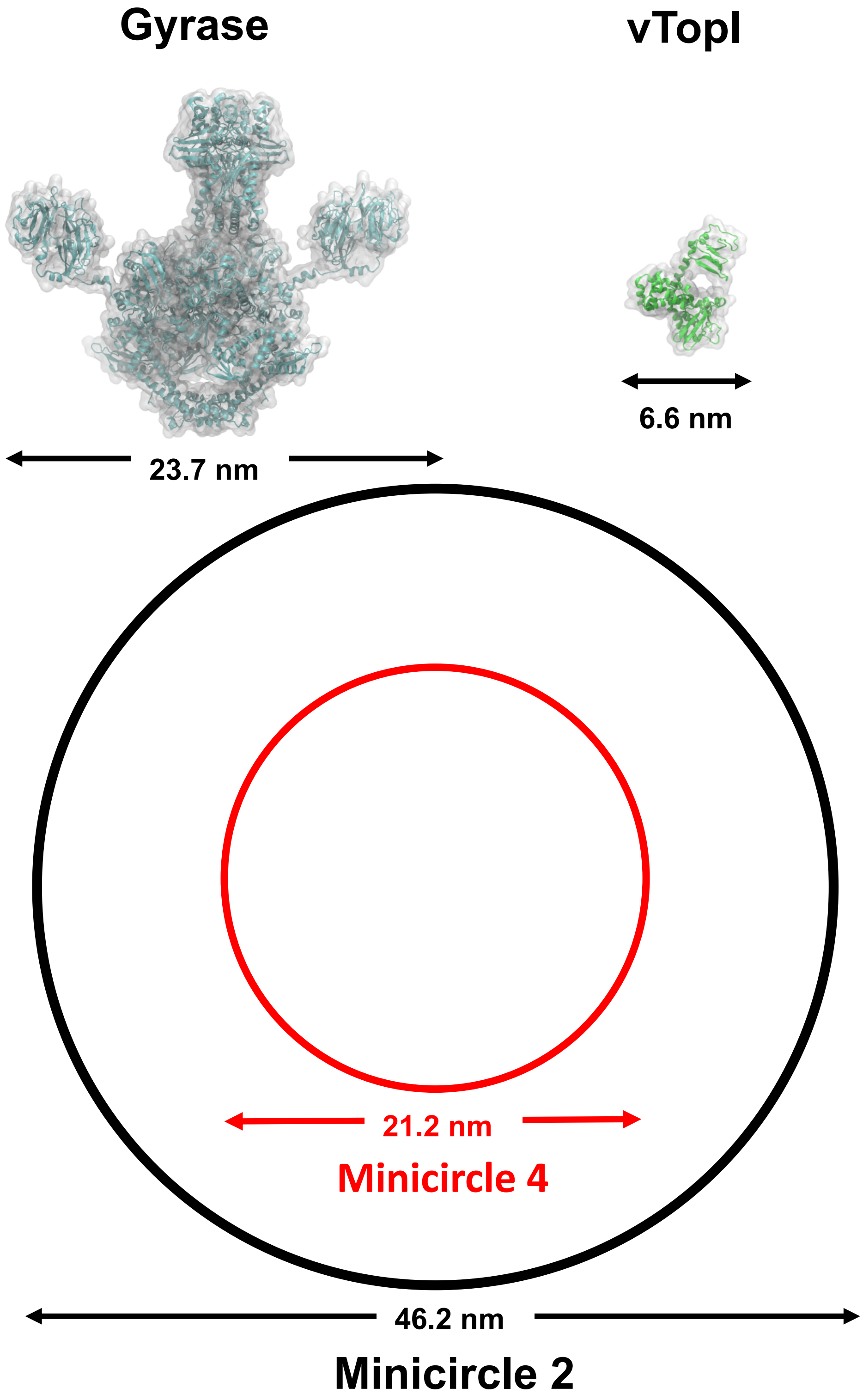


**Figure S3.** Comparing the relative sizes of *E. coli* DNA gyrase, *variola* DNA topoisomerase I (vTopI), minicircle 2, and minicircle 4.

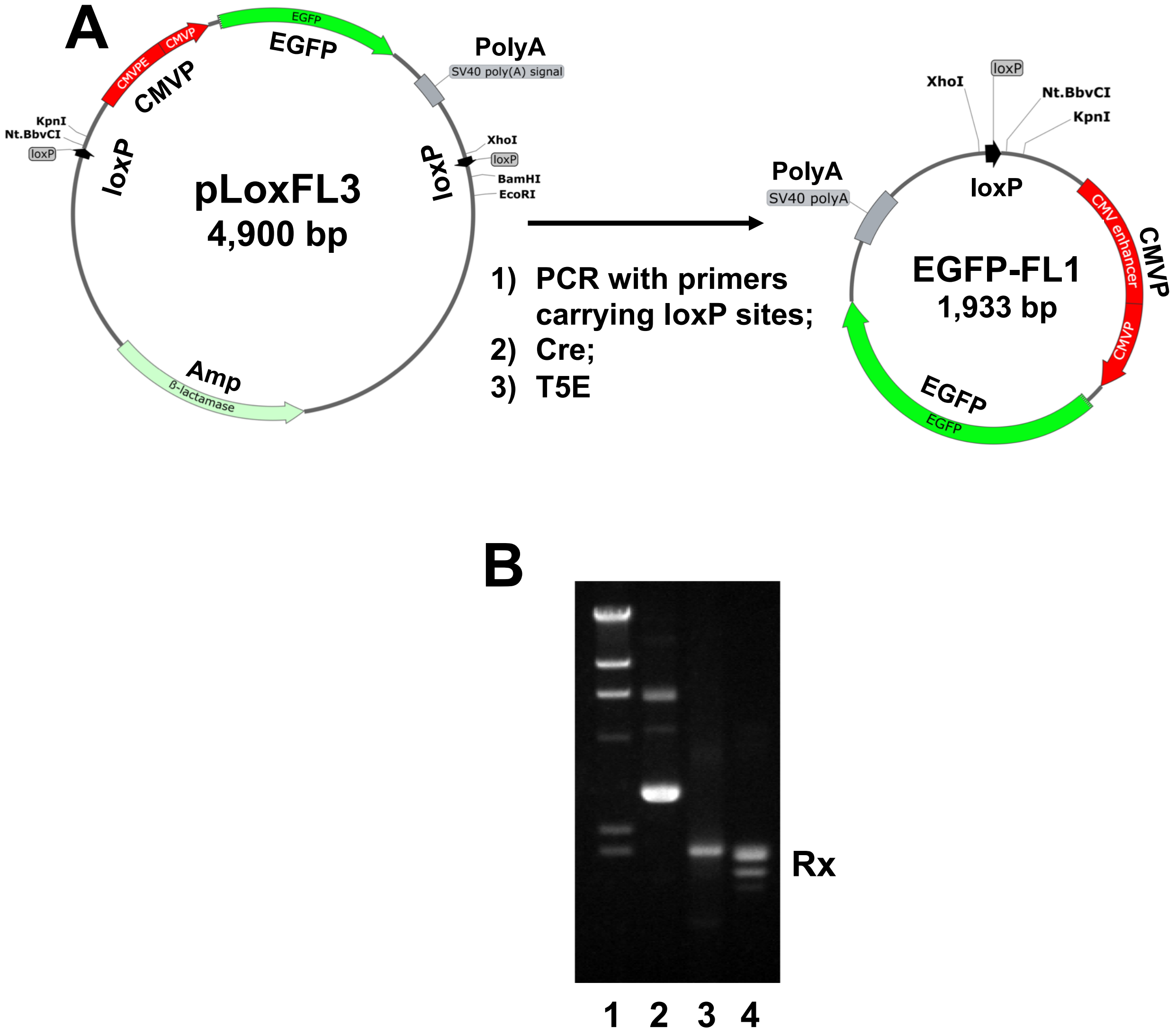


**Figure S4.** Circular DNA molecule EGFP-FL1 was produced using plasmid pEGFP-C1-FL by the PCR-based biochemical method as the DNA template that does not contain loxP sites (**A**, the procedure). Forward and reverse primers carrying a loxP site were used to amply a ??? bp DNA fragment of pEGFP-C1-FL (**B**, lane 2) to generate a linear DNA fragment carrying two loxP sites facing the same orientation (**B**, lane 1). Cre DNA recombinase converted the linear DNA molecule into Rx EGFP-FL1. T5 exonuclease was used to digest unwanted linear DNA molecules (**B**, lane 4).

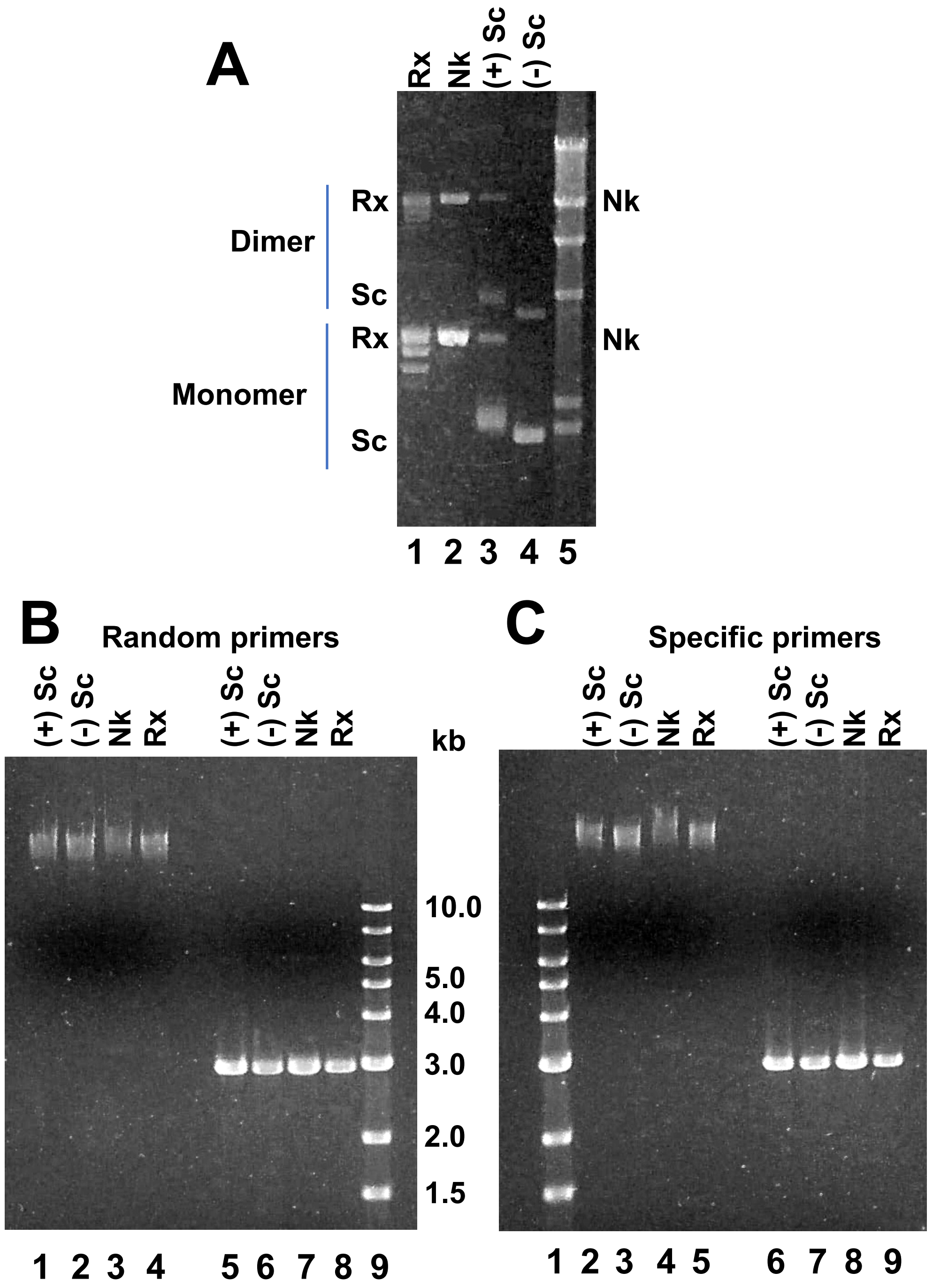


**Figure S5.** RCA of by ϕ29 DNA polymerase using different circular DNA templates, i.e., Rx, Nk, (+) Sc, and (-) Sc pLoxFL. Nk pLoxFL produced more RCA DNA products. (**A**) 1% agarose gel of Rx, Nk, (+) Sc, and (-) Sc pLoxFL. (**B**) RCA products by using Rx, Nk, (+) Sc, and (-) Sc pLoxFL as DNA templates and random hexamer primers. Lanes 1-4 are high molecular weight RCA DNA products. Lanes 5-8 are RCA products digested with BamHI. Lane 9 is $\lambda$ DNA Hind III digest. (C) RCA products using Rx, Nk, (+) Sc, and (-) Sc pLoxFL as DNA templates and two specific primers. Lanes 2-5 are high molecular weight RCA DNA products. Lanes 6-9 are RCA products digested with BamHI. Lane 1 is $\lambda$ DNA Hind III digest.

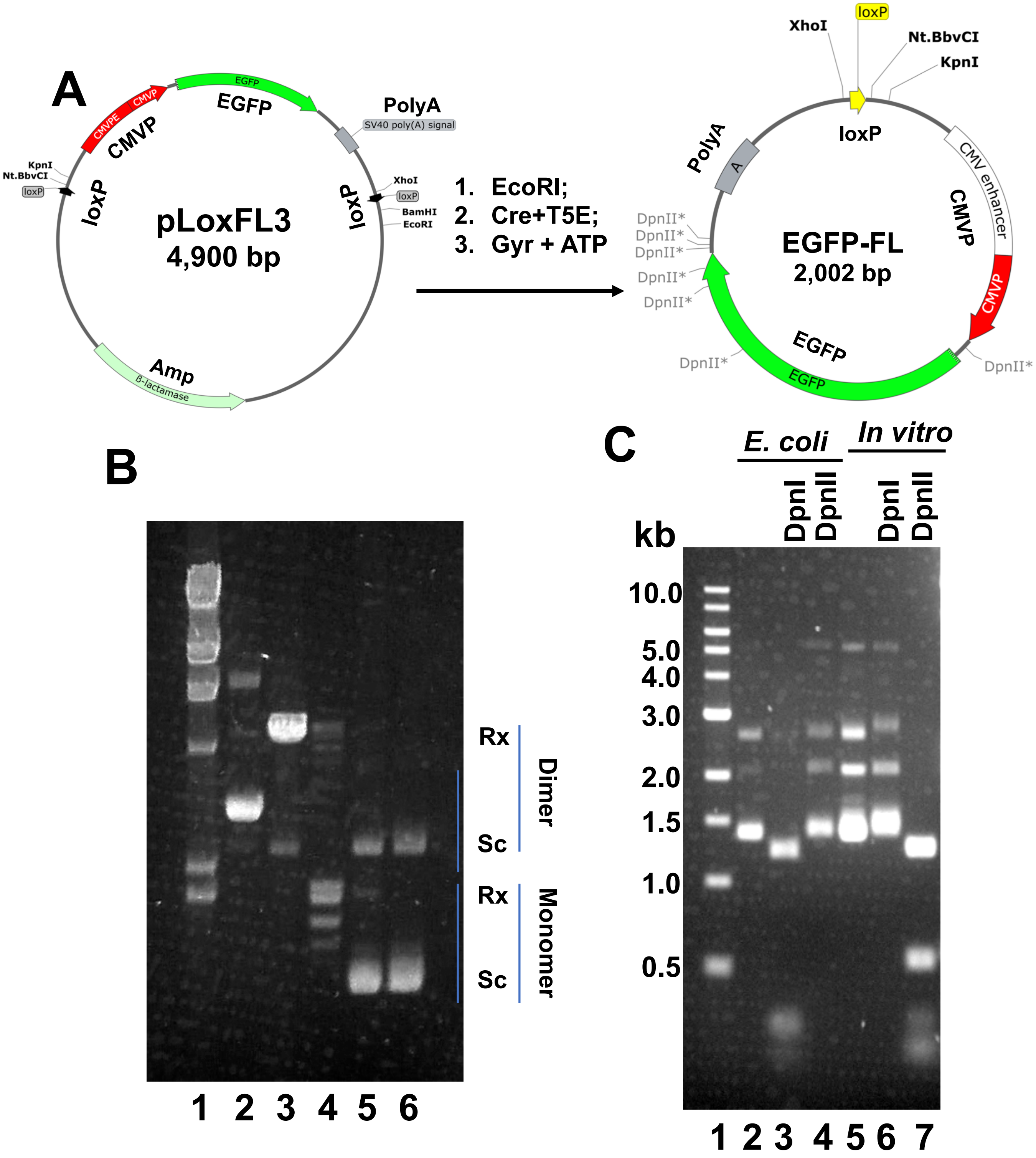


**Figure S6.** (**A**) The procedure producing the *E. coli*-derived EGFP-FL carrying Dam and Dcm methylation sites using plasmid pLoxFL3 isolated from *E. coli* cells. (**B**) 1% agarose gel showing different stages of producing the *E. coli*-derived EGFP-FL. Lane 1, λ DNA HindIII digest. Lane 2, Sc pLoxFL3. Lane 3, EcoRI-linearized pLoxFL3. Lane 4, the recombination products by Cre recombinase, digested with T5 exonuclease. Lanes 5 and 6, Sc EGFP-FL. (C) The *E. coli*-derived (*E. coli*, lanes 2-4) and the *in vitro* synthesized EGFP-FL (*in vitro,* lanes 5-7) were digested by restriction enzyme DpnI and DpnII, respectively.

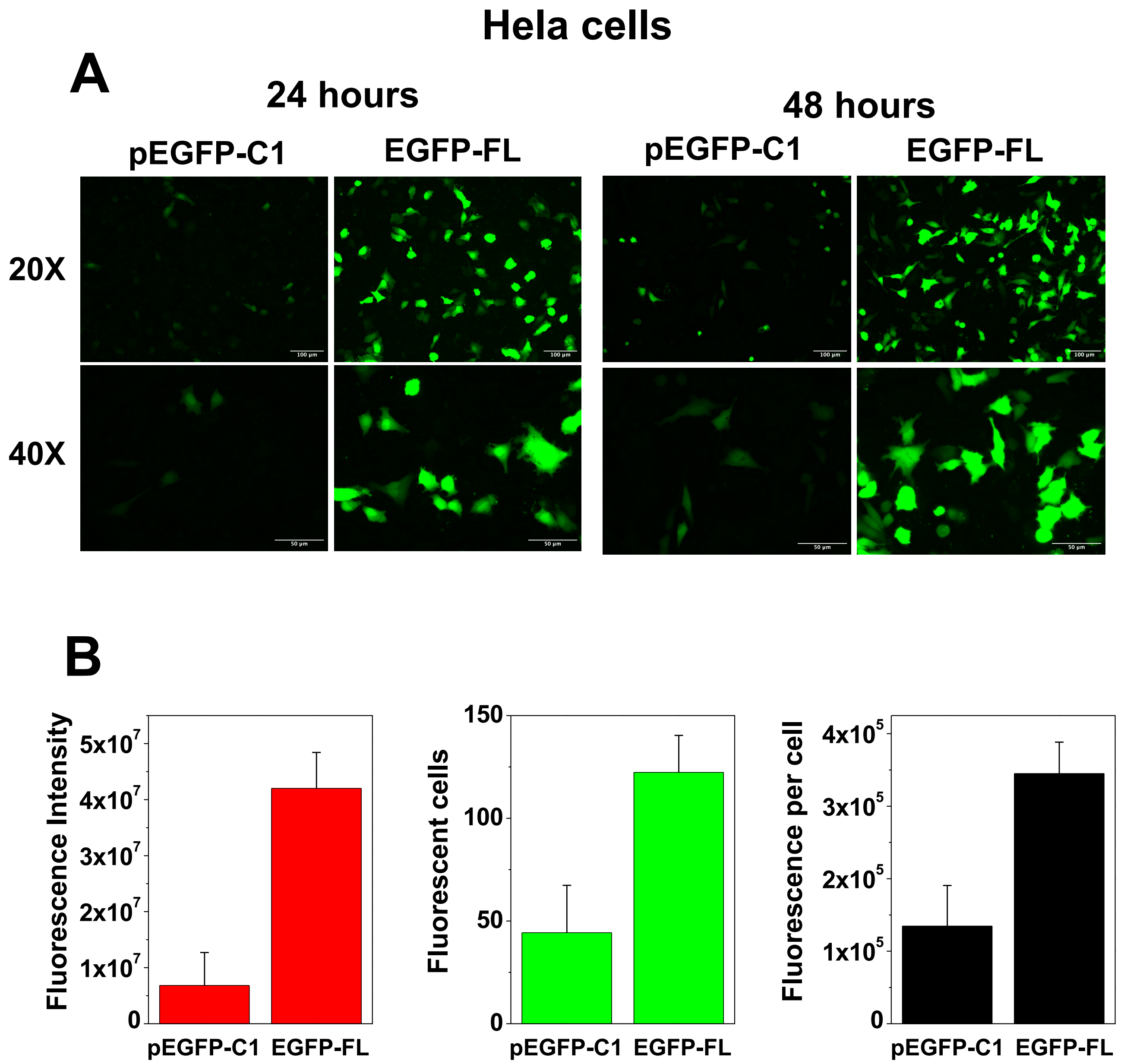


**Figure S7. Comparison of transfection efficiency of human Hela cells between the *in vitro*-synthesized, unmodified EGFP-FL and plasmid pEGFP-C1 isolated from *E. coli*.** Hela cells were seeded and grown in 96-well plates in DMEM supplemented with 10% FBS for 24 hours. Subsequently, the cells were transfected, respectively, with 0.4 μg of plasmid pEGFP-C1 isolated from *E. coli* Top10 cells or the *in vitro*-synthesized, unmodified EGFP-FL with PolyFect transfection reagent (Qiagen). 20× and 40× fluorescence images were captured at 24 and 48-hours post-transfection with a BZX800 fluorescence microscope. (**A**) Fluorescence images after 24 or 48 hours of transfection. (**B**) The fluorescence intensity of transfected Hela cells by pEGFP-C1 and the *in vitro* synthesized, unmodified EGFP-FL.

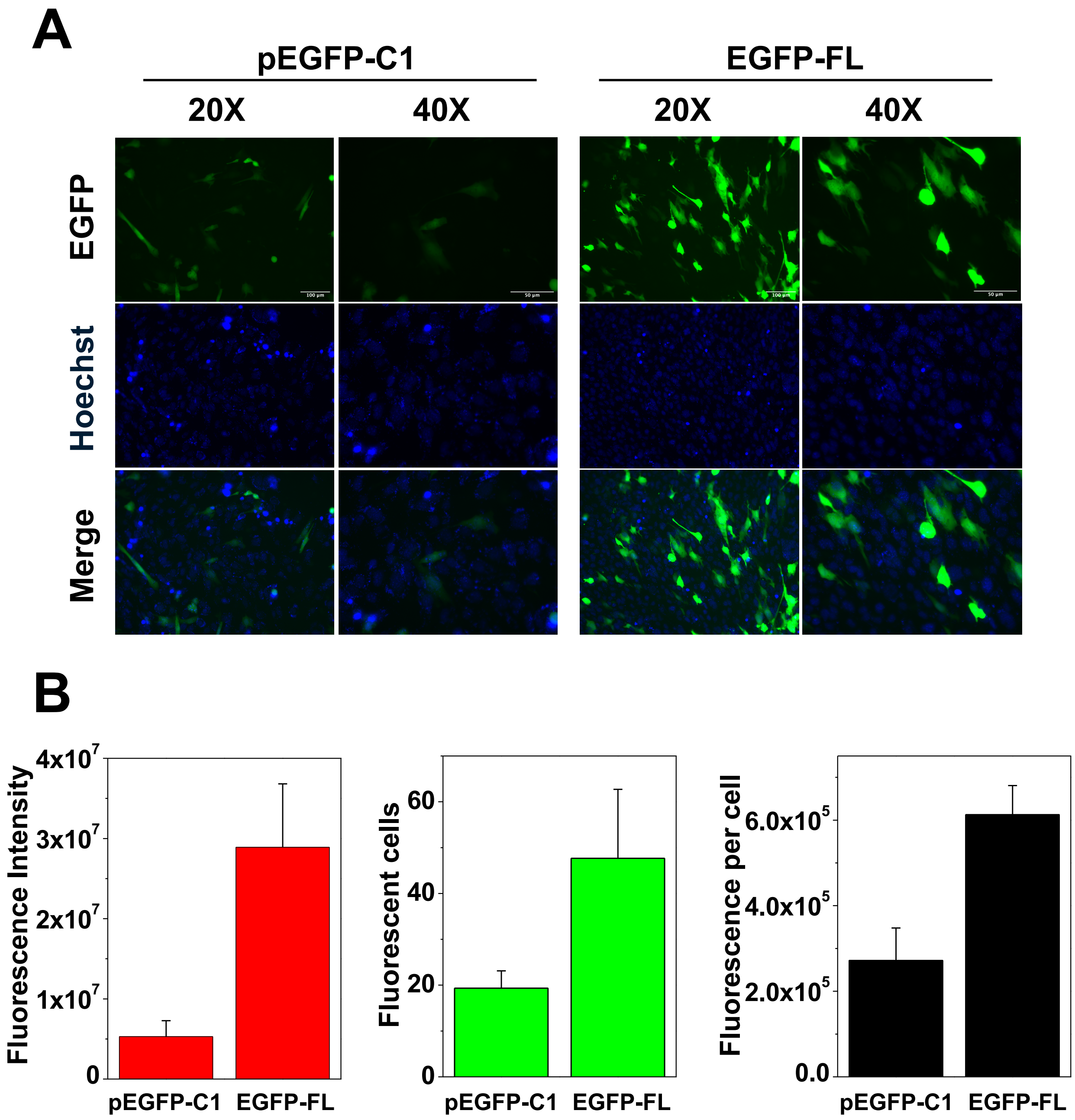


**Figure S8. Comparison of transfection efficiency of mouse myoblast C2C12 cells between the *in vitro*-synthesized, unmodified EGFP-FL and plasmid pEGFP-C1 isolated from *E. coli*.** C2C12 cells were seeded and grown in 96-well plates in DMEM supplemented with 10% FBS for 24 hours. Subsequently, the cells were transfected, respectively, with 0.4 µg of plasmid pEGFP-C1 isolated from *E. coli* Top10 cells or the in vitro-synthesized, unmodified EGFP-FL with PolyFect transfection reagent (Qiagen). 20× and 40× fluorescence images were captured at 48-hours post-transfection with a BZX800 fluorescence microscope. (**A**) Fluorescence images after 24 or 48 hours of transfection. (**B**) The fluorescence intensity of transfected Hela cells by pEGFP-C1 and the *in vitro* synthesized, unmodified EGFP-FL.

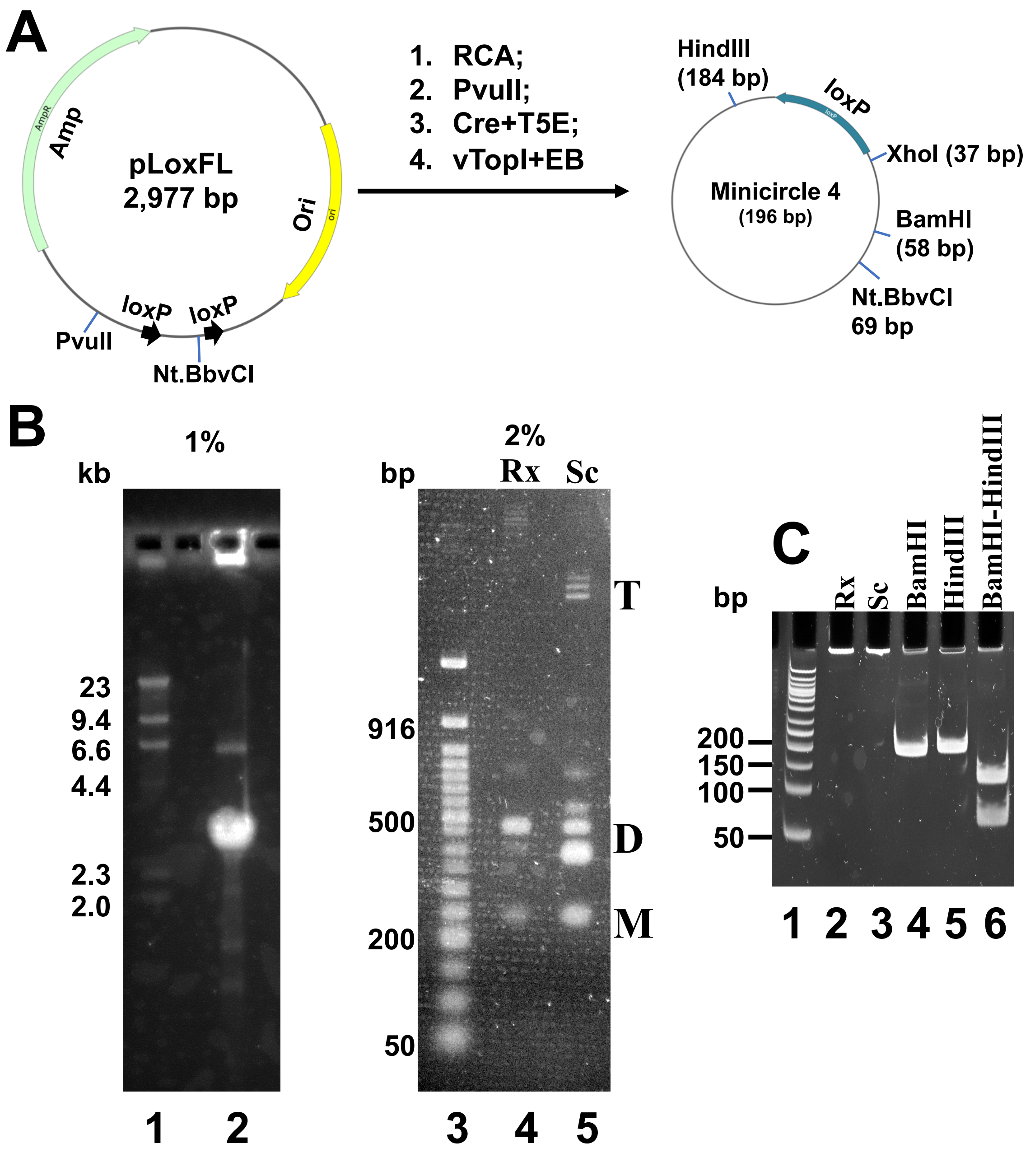


**Figure S9.** (**A**) Experimental procedure to generate the 196 bp minicircle 4 using the RCA-based biochemical method. (**B**) Left panel is a 1% agarose gel for the RCA product of Nk pLoxFL by ϕ29 DNA polymerase. Lane 1, λ DNA HindIII digest. Lane 2, the RCA product digested with PvuII. Right panel, 2% agarose gel. Lane 3, 50 bp DNA ladder. Lane 4, Rx minicircle 4. Lane 5, Sc minicircle 4. M, monomer; D, dimer; T, tetramer; Rx, relaxed; Sc, supercoiled. (**C**) Restriction digestion pattern of minicircle 4 in a 12% PAGE gel in 1×TAE buffer. Lane 1, 50 bp DNA ladder. Lane 2, Rx minicircle 4. Lane 3, Sc minicircle 4. Lanes 4 and 5, minicircle 4 digested by BamHI or HindIII, respectively. Lane 6, minicircle 4 digested by BamHI and HindIII.

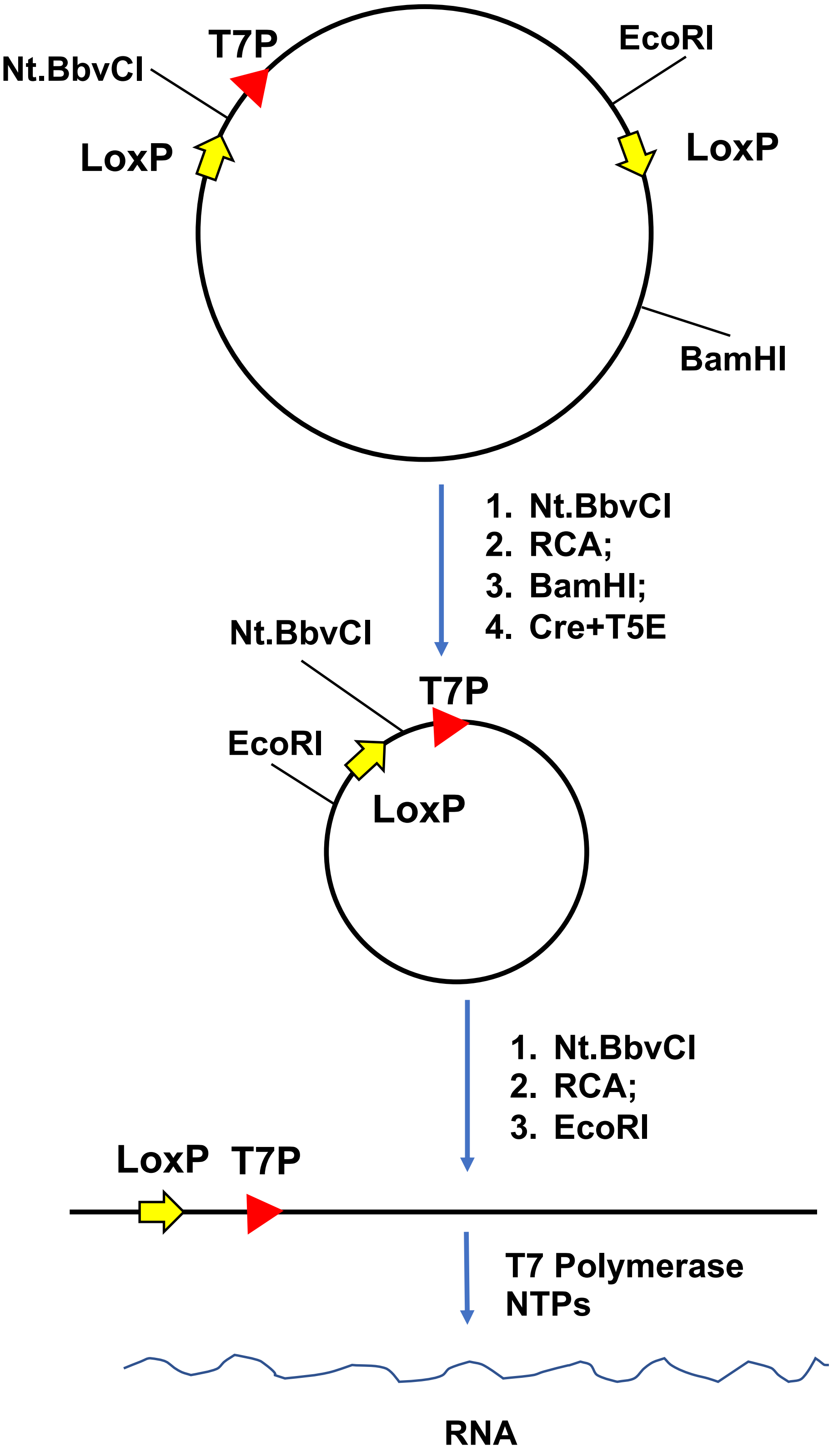


**Figure S10.** A biochemical procedure to synthesize milligrams of linear DNA for *in vitro* transcription by T7 RNA polymerase. Typically, 2 to 3 milligrams of DNA can be produced in 10 mL of reaction mixtures.
